# A robust structural, kinetic and biophysical characterization of human ACOD1 to support novel chemotherapeutic development

**DOI:** 10.1101/2025.06.13.659517

**Authors:** Brent Runge, Hande Oktay, Ian J. Fucci, Sergey G. Tarasov, Lixin Fan, Diana C. F. Monteiro

## Abstract

Human aconitate decarboxylase (ACOD1) has emerged as a therapeutic target for cancer and inflammatory diseases, with its metabolic product, itaconate, as a multifunctional metabolite shown to drive several disease states. Though extensively studied *in cellulo* and *in vivo*, this protein is biochemically and mechanistically under characterized. In this work we present a robust mechanistic study of hACOD1, with insights into its structure-activity relationships and inhibition. First true-apo and inhibited structures of this enzyme and selected point-mutants are presented, together with complementary kinetic, biochemical and biophysical analyses. We also devised a novel, low-enzyme consumption, continuous kinetic assay to support complex kinetic studies, as exemplified by the competitive inhibition assays performed. Together with molecular dynamics simulations and solution X-ray scattering experiments, all these data shed light on mechanism and dynamic behavior of the enzyme, vital insights for future drug discovery campaigns.

## 1. Introduction

Human aconitate decarboxylase 1 (hACOD1) plays a pivotal immunoregulatory role in host-pathogen interactions^1-4^, both pro- and anti-inflammatory responses^5-7^, and in the modulation of oxidative stress responses^8,9^. hACOD1 has also been specifically shown to contribute to the maintenance of tumor microenvironments^10,11^. hACOD1 is encoded by the human immune regulatory gene 1 (hIRG1) and is responsible for catalyzing the decarboxylation of *cis-*aconitate to yield the terminal metabolite itaconate in the tricarboxylic acid (TCA) cycle (Fig. 1)^12^. Itaconate was initially identified as a key metabolite highly upregulated in macrophages triggered by inflammatory stimuli, such as the recognition of pathogen-associated molecular patterns (PAMPS), including lipopolysaccharides (LPS)^13^.

**Fig. 1.**
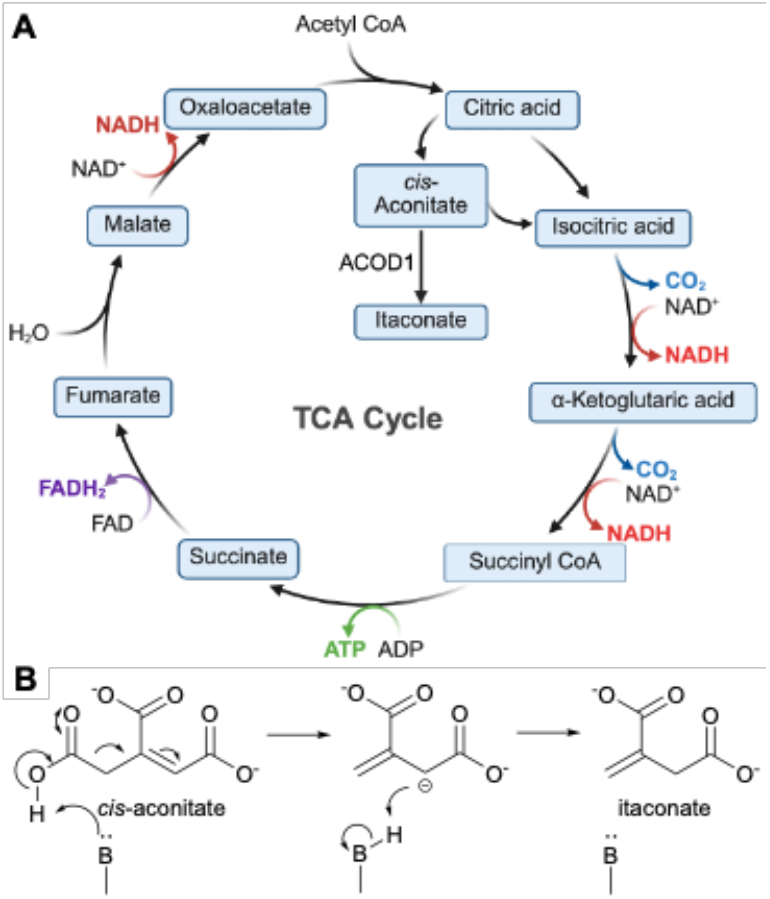
TCA cycle^14^ and hACOD1. A) overview of the TCA cycle showing metabolic intermediates, including the transformation of *cis*-aconitate to the terminal metabolite itaconate by ACOD1. B) Reaction mechanism of *cis*-aconitate decarboxylation by ACOD1.

Itaconate has a direct regulatory effect on the TCA cycle, by inhibiting succinate dehydrogenase (SDH), upregulating succinate concentration in cells, which impacts mitochondrial respiration and cytokine production in macrophages^6^. These regulatory mechanisms lead to anti-inflammatory responses including halting of mitochondrial reactive oxygen species (ROS) production and downregulation of HIF-1a activity and of IL-1b expression. Itaconate reportedly also further regulates antioxidant responses by modulation of KEAP1, stopping proteasomal degradation of the transcription factor NRF2^15^. The increase in NRF2 leads to ROS detoxification and protects cells against cytotoxic oxidative stress^16^. Itaconate also directly interacts with TET2, an important player in innate immune homeostasis and tumor suppression^17,18^. All these anti-inflammatory cascades, resulting from increased itaconate production, lead to a cellular environment more resilient to cytotoxic conditions and thus resistant to apoptotic mechanisms.

hACOD1 has been shown to have elevated expression levels in cancer cells and is a reported tumor growth factor^10,11,19,20^ making hACOD1 a promising chemotherapeutic target for sensitizing cancer cells and elicit ferroptosis^21^. Currently, citraconate is the is only reported active inhibitor of hACOD1^22^, together with some derivatives^23,24^. Citraconate is a structural mimetic itaconate (the product, Fig. 1). Though inhibition kinetics have been obtained and a proposed binding pose modelled *in silico* in the active site, no structure or full competition kinetics have been demonstrated.

In this work we fill this knowledge gap by presenting the first ligand-bound structure of hACOD1 with citraconate and first robust full inhibition kinetics showing clear competition between citraconate and *cis-*aconitate. We take the work further by determining the binding affinity of the inhibitor and the structures and biophysics of several point mutants. The mutational work indicates two important regions of the enzyme involved in its catalytic mechanism.

## 2. Results and discussion

### 2.1. An artifact-free structure of apo ACOD1

The only structure available for human ACOD1 was solved in 2019 by Chen *et. al*.^25^ together with that of the murine protein. Crystals were obtained from crystallization cocktails containing tacsimate and citrate respectively^25^, and the hACOD1 structure shows the presence of bound species in the proposed putative active site. Tacsimate is a mixture of polydentate carboxylates that are structurally similar to the substrate/product of hACOD1. It is thus, unsurprising, that these components bind to the protein, even if disorderly. A true apo crystal without promiscuous binders is necessary for clean co-crystallization with inhibitors as well as soaking methodologies for drug discovery.

After optimizing the expression and purification of hACOD1 to yield stable, active and pure protein suitable for structural and enzymatic studies (see Sup. Table 1 for final expression conditions), we set out to obtain reliable crystallization conditions for hACOD1 using high-throughput methods. Protein crystallization relies on empirical screening of crystallization conditions with chemicals known to drive crystal formation. We set 1.7k crystallization experiments, with 576 different crystallization cocktails (sparse matrix screens) and varying protein-to-cocktail volume ratios. From this initial screening campaign, we obtained ∼30 crystallization hits many of which contained unwanted polydentate carboxylates. From these, 11 conditions (Sup. Table 2) were chosen for further optimization and crystals diffracting routinely to 1.3 Å or better were obtained from two of the initial hits. Upon structure solution from data collected from crystals grown in sodium acetate, no evidence of bound acetate was found in the active site, showing only well-ordered water molecules, and indicating that our high-throughput crystallization pipeline was successful in producing a true apo hACOD1 structure at high resolution of 1.22 Å (Fig. 2, B, Table 1).

**Table 1.** X-ray data and model statistics.

|  | apo hACOD1 | citraconate-hACOD1 | hACOD1-R273S | hACOD1-Y318A | hACOD1-H103A |
| --- | --- | --- | --- | --- | --- |
| PDB ID | 12UX | 12VD | 12VW | 12VT | 12VQ |
| Raw Data DOI |  |  |  |  |  |
| Diffraction Source | NSLSII 17-ID-1 (AMX) | NSLSII 17-ID-2 (FMX) | NSLSII 17-ID-1 (AMX) | NSLSII 17-ID-1 (AMX) | NSLSII 17-ID-1 (AMX) |
| Wavelength (Å) | 0.920 | 0.979 | 0.919 | 0.920 | 0.920 |
| Temperature (K) | 100 | 100 | 100 | 100 | 100 |
| Detector | EIGER 9M | EIGER 16M | EIGER 9M | EIGER 9M | EIGER 9M |
| Crystal to detector distance (mm) | 161.33 | 158.91 | 161.33 | 115.69 | 115.69 |
| Total rotation range (°) | 360° | 360° | 360° | 360° | 360° |
| Rotation per image (°) | 0.2° | 0.2° | 0.2° | 0.2° | 0.2° |
| Exposure time per images | 5 ms | 5 ms | 5 ms | 5 ms | 5 ms |
| Space group | P2<sub>1</sub>2<sub>1</sub>2 | P2<sub>1</sub>2<sub>1</sub>2 | P2<sub>1</sub>2<sub>1</sub>2 | P2<sub>1</sub>2<sub>1</sub>2 | P2<sub>1</sub>2<sub>1</sub>2 |
| a, b, c (Å) | 101.8, 110.1, 75.8 | 102.2, 110.4, 76.2 | 101.87, 110.19, 75.73 | 101.71, 109.99, 76.01 | 101.85, 110.04, 75.87 |
| α β, γ (°) | 90.0, 90.0, 90.0 | 90.0, 90.0, 90.0 | 90.0, 90.0, 90.0 | 90.0, 90.0, 90.0 | 90.06, 90.06, 90.08 |
| Resolution range (Å) | 33.78-1.35 (1.49-1.32) | 33.9-1.22 (1.45-1.22) | 44.55-1.46 (1.54-1.46) | 34.55-1.62 (1.817-1.62) | 34.50-1.42 (1.54-1.42) |
| Ellipsoid diffraction limits (a*, b*, c*): | 1.39, 1.43, 1.51 | 1.32, 1.44, 1.51 | 1.47, 1.48, 1.59 | 1.78, 1.70, 1.82 | 1.496, 1.489, 1.534 |
| Total no. of reflections | 2000190 (84956) | 2117439 (93179) | 1128351 (62100) | 1120717 (57300) | 1805157 (92007) |
| No. of unique reflections | 146042 (7306) | 155814 (7790) | 129053 (6453) | 80794 (4040) | 129811 (6491) |
| Completeness (%) | Spherical: 74.0 (13.0) Ellipsoidal: 94.1 (55.6) | Spherical: 61.4 (7.8) Ellipsoidal: 94.7 (54.9) | Spherical: 87.1 (28.4) Ellipsoidal: 96.2 (64.3) | Spherical: 74.0 (12.9) Ellipsoidal: 94.4 (46.3) | Spherical: 80.4 (17.6) Ellipsoidal: 95.4 (51.9) |
| Redundancy | 13.7 (11.6) | 13.6(12.0) | 8.7 (9.6) | 13.9 (14.2) | 13.9 (14.2) |
| <I/σ(I)> merged data | 9.8 (1.0) | 10.3 (1.7) | 8.4 (1.5) | 8.4 (1.5) | 10.1 (1.4) |
| CC1/2 | 0.998 (0.331) | 0.996 (0.415) | 0.997 (0.621) | 0.995 (0.550) | 0.998 (0.598) |
| Rmerge | 0.204 (2.503) | 0.187 (1.98) | 0.136 (1.48) | 0.260 (2.35) | 0.176 (2.43) |
| Rmeas | 0.220 (2.742) | 0.202 (2.15) | 0.154 (1.65) | 0.28 (2.52) | 0.189 (2.62) |
| Rpim | 0.057 (0.789) | 0.053 (0.595) | 0.050 (0.537) | 0.072 (0.664) | 0.049 (0.696) |
| Wilson B factor (Å<sup>2</sup>) | 11.4 | 9.9 | 17.5 | 19.2 | 16.1 |
| Refinement |  |  |  |  |  |
| Resolution range (Å) | 33.78-1.32 (1.48-1.32) | 33.9-1.22 (1.45-1.22) | 44.55-1.46 (1.54-1.46) | 34.55-1.62 (1.82-1.62) | 34.53-1.42 (1.55-1.42) |
| No. of reflections, working set | 146042 | 155813 | 129053 | 80793 | 129811 |
| No. of reflections, test set | 7369 | 8014 | 6449 | 4198 | 6630 |
| Final Rcryst | 15.8 | 15.1 | 0.154 | 0.16 | 0.15 |
| Final Rfree | 19.3 | 17.9 | 0.184 | 0.202 | 0.181 |
| No. of non-H atoms | 15462 | 14434 | 15172 | 15001 | 15516 |
| Protein/nucleic acid | 14417 | 14434 | 14179 | 14141 | 14433 |
| Ions | 2 | 2 | 1 | 4 | 4 |
| Ligands | 37 | 28 | 19 | 15 | 37 |
| Waters | 1006 | 1026 | 973 | 841 | 1042 |
| R.m.s deviations |  |  |  |  |  |
| Bonds (Å) | 0.01 | 0.01 | 0.0093 | 0.007 | 0.0092 |
| Angles (°) | 1.823 | 1.896 | 1.729 | 1.538 | 1.73 |
| Average B factors (Å<sup>2</sup>) |  |  |  |  |  |
| Protein/nucleic acid | 16.3 | 15 | 18.9 | 19.4 | 16.2 |
| Ions | 26.5 | 20.6 | 27.5 | 23 | 14.9 |
| Ligands | 46.9 | 19.9 | 31.5 | 35.8 | 29.1 |
| Waters | 29.7 | 28 | 30.3 | 27.4 | 28.3 |
| Ramachandran plot (Refmac5) |  |  |  |  |  |
| Favored regions (%) | 95.31 | 95.53 | 95.3 | 95.2 | 95.6 |
| Outliers (%) | 0.22 | 0.22 | 3.9 | 4 | 3.7 |
| Unmodelled/incomplete residues (%) | 0 | 0 | 0 | 0 | 0 |

**Fig. 2.**
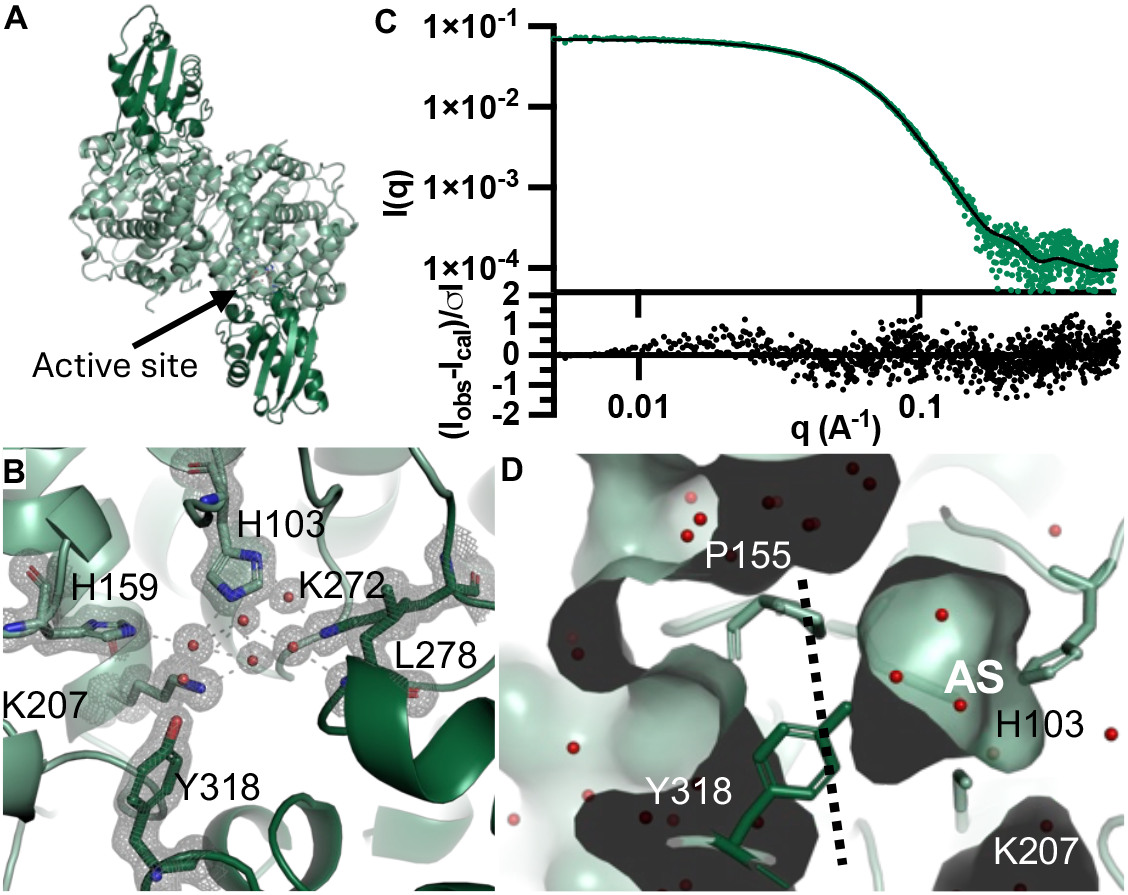
hACOD1 apo structure, active site cavity and hydrogen bonding network. A) overall architecture of the hACOD1 dimer, showing the lid domain in dark green and the alpha-helical domain in light green. B) The active site of WT hACOD1, showing ordered waters (red spheres) and the hydrogen bonding network (black dashed lines). C) Experimental SAXS profile of WT hACOD1 (green dots) fitted to the calculated solution scattering profile derived from the X-ray crystal structure (black line), showing good agreement with low weighted fit residuals (black dots). D) Surface and cavities representation of hACOD1 apo structure, showing the small active site cavity located between the two domains. The cavity is inaccessible to the bulk solvent (black dashed line shows the boundary formed by Tyr318 and Pro155).

hACOD1 is a dimer, with each protomer composed of two well-folded domains, a lid domain composed of residues 273-410 and an alpha-helical domain, residues 1-267 and 413-461 (Fig. 2, A), matching the typical topology of the MmgE/PrpD superfamily (EMBL-EBI InterPro). The catalytic sites are located between the two domains, with catalytically relevant residues His103, His159 and Lys 272 in the alpha-helical domain and Tyr318 in the lid domain (Fig. 2, B). The small putative proposed active site in the apo hACOD1 crystal structure is closed to the outside medium, through a hydrophobic barrier at the entrance by Try318 and Pro155 (Fig. 2, D). To verify that the crystallographic structure matches the native structure in solution, we collected Small-Angle X-ray Scattering (SAXS) data (Fig. 2, C). The SAXS data showed a single, well-folded species in solution and fitting to the calculated scattering curve from our experimental X-ray structure shows very good agreement. This correlation shows that this two-domain protein adopts a closed conformation in solution, rather than explore large conformational changes.

### 2.2. Structural comparison of apo-from and citraconate-bound ACOD1

The inhibitor-bound structure was obtained from co-crystallization of hACOD1 in the presence of citraconate. Crystals for the complex were grown in similar conditions to that of the apo protein (Sup. Table 3) and diffracted to high resolution (1.32 Å, Table 1). Fig. 2 B and Fig. 3 A/B show the active site architectures of apo- and citraconate-bound hACOD1 (see Fig. 3, C for overlay). The apo structure shows six highly ordered water molecules, making up a hydrogen-bonding network linking residues His103, His159, Lys272, Lys207 and Tyr308. Citraconate binding displaces 5 of the waters with the oxygen atoms taking the positions of the displaced waters and thus satisfying most of the hydrogen-bonding network. The citraconate-bound structure suggests that the ligand binds at the active site and corroborates the putative active site proposed by Chen *et al*^25^. To pursue robust kinetic characterization of hACOD1 and unequivocally prove competitive inhibition with substrate (since a substrate- or product-bound structure could not be obtained), we developed a robust new kinetic assay, described in the next section.

**Fig. 3.**
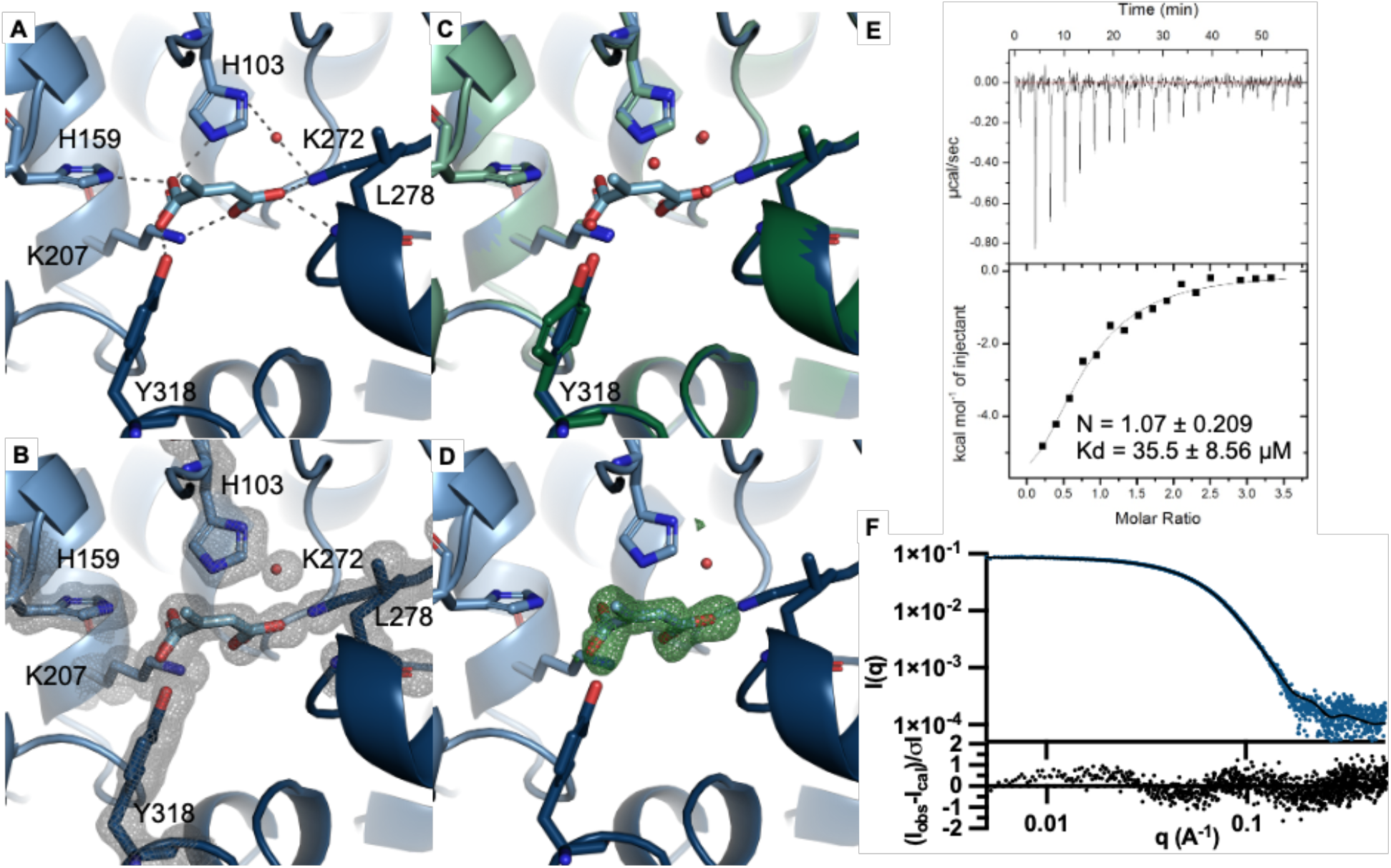
Crystal structures of apo hIRG1 (green) and citraconate-bound hACOD1 (blue). A) Citraconate-bound hACOD1 showing the hydrogen-bonding network between the inhibitor and active site residues and B) the 2*F*_o_-*F*_c_ electron density map contoured at 1σ. C) Overlay of apo and citraconate-bound structures, showing minimal movement of residues in the active site, except for Tyr318, which is partly rotated. The waters in the apo form sit in positions occupied by oxygens of the citraconate carboxylates. D) Citraconate-removed *F*_o_-*F*_c_ omit map, showing clear density for the ligand contoured at 3σ. (E) ITC titration curve of citraconate into hACOD1. (F) SAXS scattering profile of citraconate-hACOD1 (blue dots) fitted to the calculated solution scattering profile (black line) derived from the hACOD1–citraconate X-ray crystal structure (with weighted residuals).

The root mean square deviation (RMSD) between the apo and citraconate-bound structures is very low (0.097 Å globally and 0.108 Å between same chains), indicating that the protein overall conformation does not differ between the apo and ligated forms. As with apo hACOD1, the citraconate-bound structure is a compact dimer in solution (Fig. 3, F). From all the residues comprising the active site and part of hydrogen-bonding network, only Tyr318 shows a conformational change between the two structures, and it is minimal at 1.7 Å (Fig. 3, C). hACOD1 follows a typical acid-base catalytic mechanism^26^, but the identity of the base(s) involved in the proton transfer mechanism is still unknown. The active site, now confirmed by our citraconate-bound structure and competitive inhibition kinetics (see next section), is well-defined and previous work proposes that His103 should be involved due to its proximity^25^. In our ligand-bound structure, His103 is indeed well position abstract a proton at the C2 position of citraconate.

### 2.3. Development of a low-consumption, continuous enzymatic assay for hACOD1

To obtain robust kinetic parameters for activity and inhibition by hACOD1, we developed a novel ^1^H NMR-based catalytic assay. This assay has several advantages over the previous reported end-point HPLC assay^25^. First, it allows for the direct visualization of reaction progress and quantification of product formation/substrate consumption by integration of assigned proton signals. Second, it is a continuous assay, where multiple time-points can be measured from the same sample, greatly diminishing sample consumption and reducing experimental error from sample preparation. Third, the continuous assay allows for the determination of initial vs steady state rates in one measurement, and robust determination of initial velocities from multi-point fitting. By interleaving data collection of multiple samples and measuring multiple time-points from a single sample, full kinetics can be obtained in triplicate within 48h of machine time and using only 15 μg of protein per substrate/inhibitor concentration.

In this assay, both the product and substrate signals can be concurrently measured as they have distinct chemical shifts. Decarboxylation of *cis*-aconitate by hACOD1^26^ converts its tri-substituted alkene to a geminal alkene (Fig. 4, A). The signals for the protons on the alkene are distinct and quantifiable ^1^H NMR (Fig. 4, B, Sup. Fig. 1). Quantification can be done either by the ratio of proton integrals for the substrate and product or direct quantification using an internal standard. The initial rate for each kinetic assay was derived by fitting a linear regression to the linear regime of itaconate production, corresponding to the initial ∼10-20% turnover of *cis*-aconitate. Given the temperature-dependence of ^1^H chemical shifts, we found that temperature equilibration of samples prior to adding protein to solution was necessary to extract accurate measurements. hACOD1 catalytic properties were modelled using Michaelis-Menten kinetics, giving a k_cat_=0.83±0.03 s^-1^ and K_M_=0.57±0.06 mM. These values were comparable to those previously reported.

**Fig. 4.**
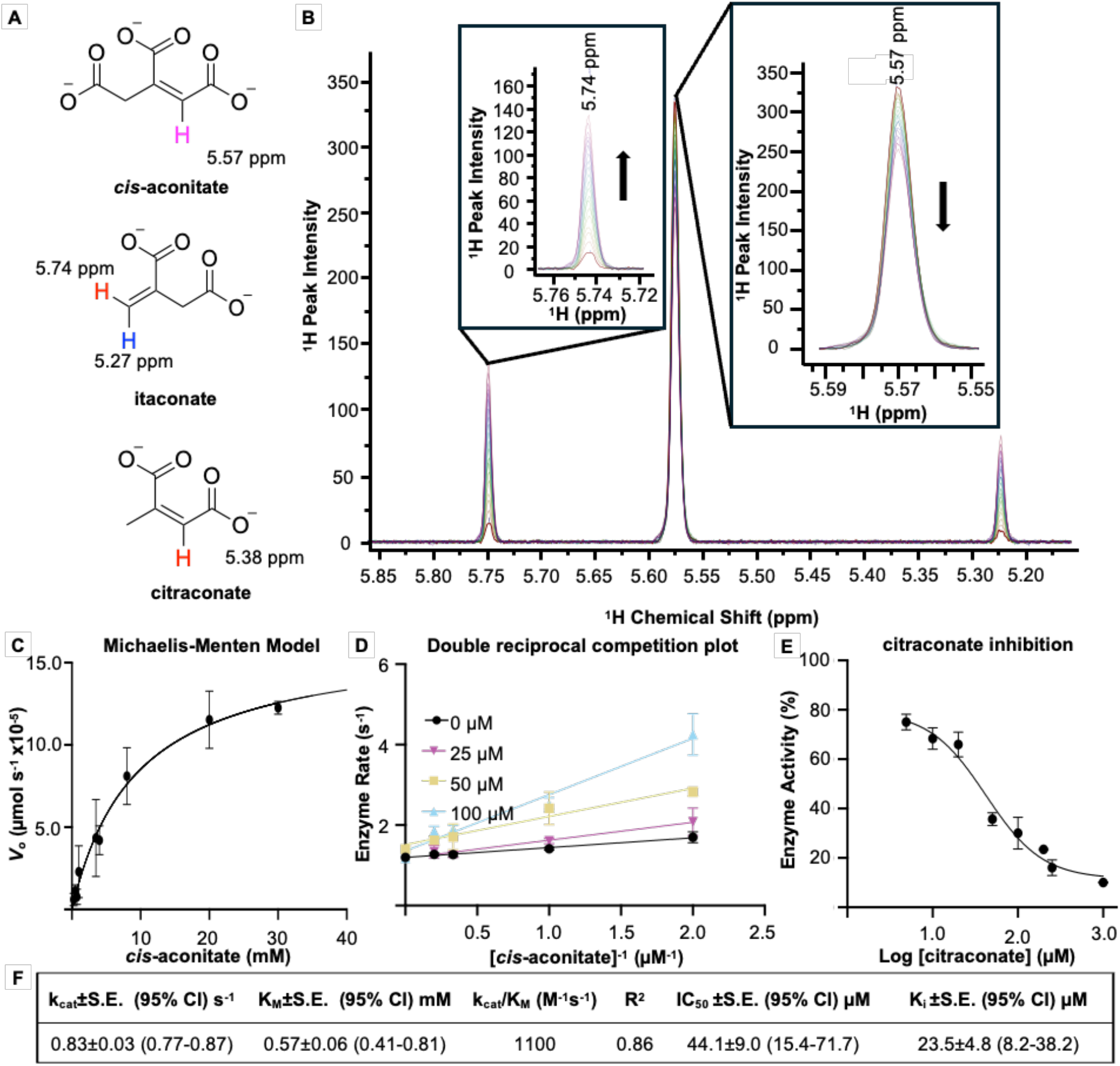
Kinetics and inhibition of hACOD1 assays by ^1^H NMR. A) structures of *cis*-aconitate, itaconate and citraconate showing the distinct chemical shifts associated with the alkene protons for each species. B) Overlay of ^1^H NMR spectra of hACOD1 kinetic assay over a two-hour time course. C) hACOD1 enzymatic activity characterization by Michaelis-Menten kinetics. D) Double reciprocal plot of hACOD1 inhibition by citraconate showing clear direct competition with substrate. E) Dose-dependent inhibition of hACOD1 by citraconate for IC5_0_ calculation. F) Kinetic and inhibition constants for hACOD1.

We used this newly developed assay to determine the inhibition kinetics of hACOD1 by citraconate. While similar in structure, citraconate has distinct proton chemical shifts from *cis*-aconitate and itaconate, and thus the same data analysis methodology could be used (Fig. 4, A, Sup. Fig. 1). Similarly to the assays preformed on apo-protein, two-hour time-course experiments were used to measure reduction in hACOD1 enzymatic activity in the presence of inhibitor. The *in vitro* IC_50_ measured in this study of 44.1 µM is comparable to that previously determined by cellular studies (Fig. 4, E/F)^22^. To show the useability of our new, low consumption assay, and provide unequivocal evidence that citraconate is a competitive inhibitor and obtain a K_i_, we expanded our kinetic assay to obtain a Lineweaver-Burk plot. The y-axis intercept at 1/Vmax with changing citraconate concentration together with changing of x-axis intercept (not shown) shows clear competitive inhibition yielding a final K_i_ of 23.5 µM.

And finally, for full characterization of this inhibitor, we also measured the binding thermodynamics of the hACOD1-citraconate interaction using isothermal titration calorimetry (ITC, Fig. 3, E). Fitting of the ITC data to a single site mechanism yields a K_d_ of 35.5 μM, within range of the calculated IC_50_ for this inhibitor. The binding is strongly enthalpic (ΔH=11 kcal•mol^-1^) and shows a 1:1 interaction between protomer and ligand (2 ligand molecules per hACOD1 dimer).

### 2.4. Mutagenesis and molecular dynamics studies shed light on active site mechanisms

With a robust kinetic assay and thermodynamic measurements in hand together with high-resolution, artifact-free crystal structures of the apo and inhibited protein, we set out to do further mechanistic studies. From a sequence alignment of hACOD1 homologues and comparison to our crystal structures, we chose 4 mutants aimed at determining structure-activity relationships.

The first mutant investigated was hACOD1-His103Ala, where a fully conserved residue in the active site was mutated. From the ligated structure and proposed mechanism, His103 is known to be involved in the catalytic mechanism, as His103 mutants showed no appreciable activity in previous work^25^. The point mutant yielded stable, well folded protein (Fig. 5, A, Sup. Fig. 2) and, as expected, with no visible catalytic turnover in our assay. The crystal structure shows a well-defined active site, matching the original apo structure, but with changes to the ordered water network as the H-bond donor/acceptor pair of the histidine residue is removed. Furthermore, the structure also shows the presence of a bound glycerol molecule in the active site, promoted by the enlarged active site cavity from the reduced Ala103 amino-acid size. To elucidate whether His103 is only necessary for catalysis or also affects binding, we performed an ITC binding experiment with citraconate, which showed no measurable binding. Co-crystallization in the presence of citraconate showed no evidence of binding from electron density (not shown). His103 is therefore necessary at the binding stage of the catalytic cycle.

**Fig. 5.**
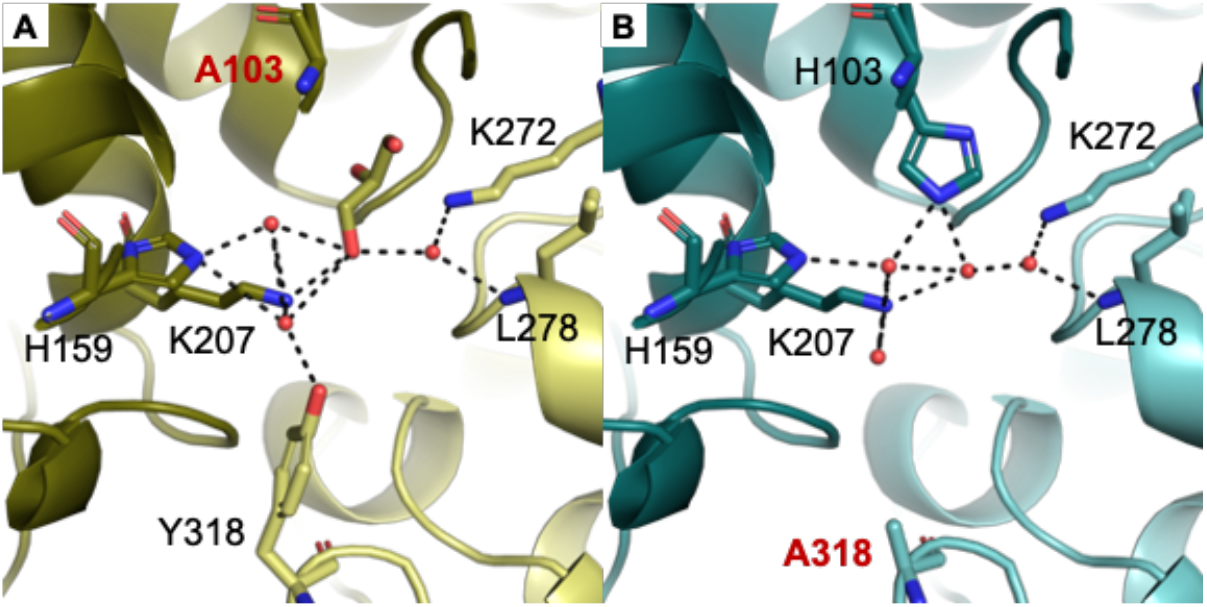
hACOD1-His103Ala and -Tyr318Ala mutants X-ray structures in yellow and teal (A and B respectively), showing the hydrogen bonding network within the active site. The His103Ala mutation opens the active site cavity and allows for the accommodation of a buffer glycerol molecule.

The WT apo hACOD1 structure shows that the active site is small and shielded from bulk solvent by loops Asp153-His159 Ile313-Gln319, specifically with Pro155 and Tyr318 blocking the active site entrance (Fig. 2, D). We thus proposed that Pro155 and Tyr318 should be important for hACOD1 function, where Pro155 is not a conserved residue and Tyr318 is semi-conserved.

The hACOD1-Pro155Ala mutant was found to undergo slow aggregation after purification, preventing crystallization, and thus a structure could not be obtained. With freshly prepared protein, kinetic assays showed no catalytic activity and ITC no binding to citraconate. Pro155, though not conserved, plays an important part on protein stability and function. By contrast, hACOD1-Tyr318Ala yielded stable, well folded protein (Fig. 5, B and Sup. Fig. 2), but not catalytically active and unable to bind citraconate. Inspection of the hACOD1-Tyr318Ala structure (Fig. 5, B) shows a well-ordered active site, with fewer water molecules visible compared to the WT protein, indicating a loss of some of the hydrogen bonding network. The active site cavity is now exposed to the bulk solvent (Fig. 6, D). hACOD1 homologues show phenylalanine as an alternative residue at this position: a large, hydrophobic residue, with no hydrogen-bond forming sidechain. From these observations, stable binding of molecules to in the active site requires the shielding of the cavity but not a hydrogen bond donor at this position, as initially expected from the hACOD1-citraconate structure where the hydroxyl group of Ty318-OH interacts with the ligand. For catalysis to occur, Tyr318 must change conformation to allow for access to the binding site cavity during substrate binding and product release.

**Fig. 6.**
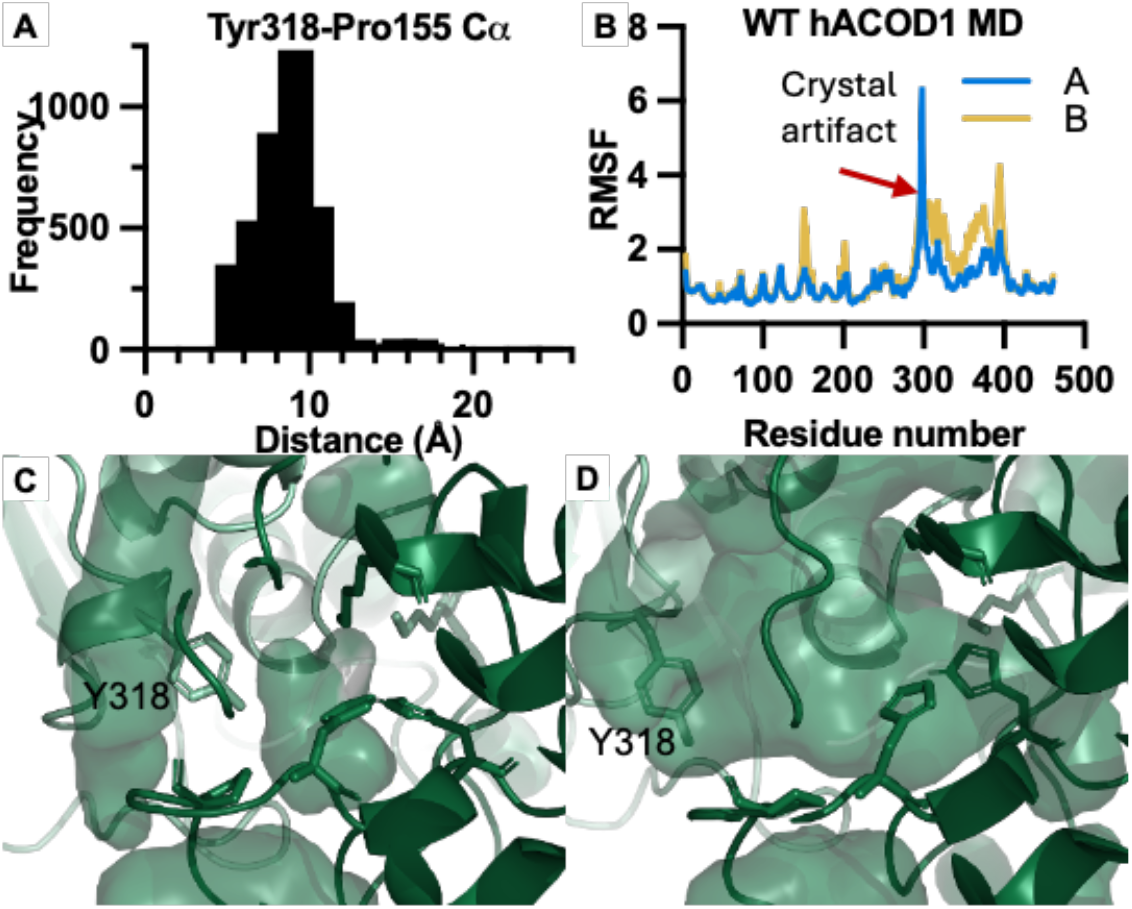
Dynamics in hACOD1. A) and B) MD simulation results for WT apo hACOD1, showing the tight distance distribution between Tyr318 and Pro155 Cα (A) and the general dynamics across the whole protein as a root-mean-square-fluctuation (RMSF, B). The RMSF shows similar dynamic patterns across the two protomers, with differences between the chains propagated from the initial crystallographic structure. C) and D) The active site cavity and accessibility of the closed and open conformations, shown by surface representation. Swinging of Tyr318 causes the small, shielded active site cavity to open to open to the bulk solvent.

**Fig. 7.**
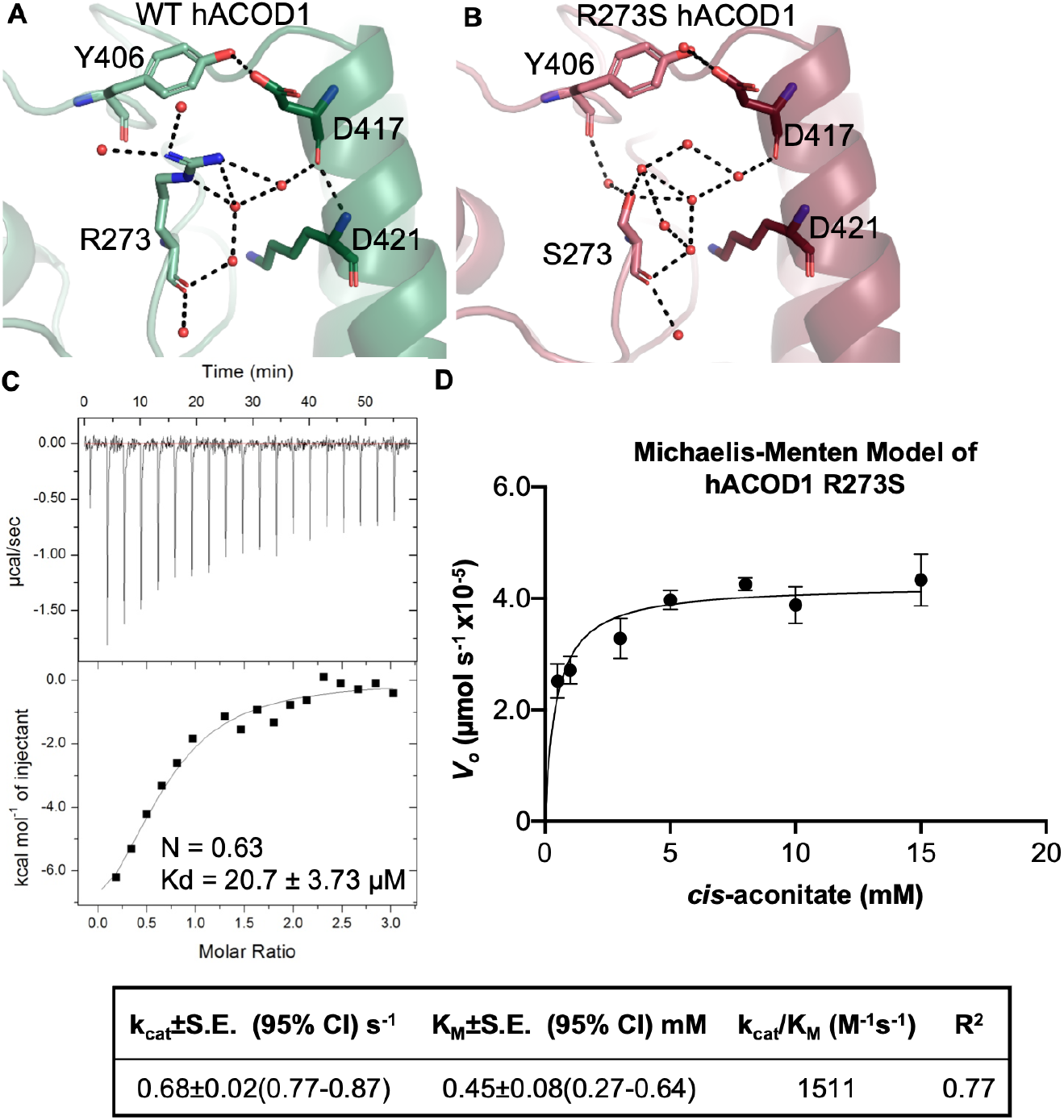
hACOD1-Arg273Ser mutant characterization. A) and B) show the local structure and water network surrounding residue 273, which is solvent exposed. The single-water bridge between Arg273 and Asp417 is lost upon mutation to serine. C) ITC of hACOD1-Arg273Ser mutant binding to citraconate. D) Michaelis-Menten kinetic model of hACOD1-Arg273Ser.

To further elucidate the dynamics involved in this mechanism, we ran Molecular Dynamic (MD) simulations starting from our WT apo hACOD1 X-ray structure and analyzed our SAXS data further. Molecular dynamic simulations of the dimeric hACOD1 show clear conformational fluctuations around Tyr318 and Pro155 (in the alpha-helical and lid domains respectively). Plotting the mean distance between the alpha-carbons of these two residues samples during the simulation shows that their relative positions fluctuate minimally at around 9 Å (Fig. 6, A). SAXS data for the WT and Tyr318Ala mutant fit well to the solved crystal structures (Fig. 2, C and Sup. Fig. 2), corroborating that the global conformation of these proteins in solution is similar to that obtained by crystallography. To assess flexibility, we performed a dimensionless Kratky Analysis (Sup. Fig. 2), which showed low flexibility, similar across all the datasets, including the citraconate-bound WT hACOD1 sample. Together, these observations indicate that necessary conformational changes associated with accessibility of the active site occur on a local scale, rather than a full domain open/closing mechanism. Frames from the MD trajectory indeed show different possible conformations for Tyr318, where the active site becomes accessible from swinging of the tyrosine side chain (Fig. 6, D).

### 2.5. Molecular dynamics and mutagenesis studies shed light on allosteric mechanisms

Further inspection of frames from the MD trajectory, revealed significant local dynamics and conformational sampling. We decided to further investigate one such mobile residue, Arg273, a solvent exposed residue next to the conserved active-site Lys272. In murine ACOD1, residue 273 is a serine and the enzyme is known to be faster and more efficient. Previous work showed that an Arg273His mutation (not naturally occurring to our knowledge) leads to a decrease in K_M_, with no significant change in k_cat_.

hACOD1-Arg273Ser indeed also shows changes in kinetic properties, with a slight decrease in K_M_ and k_cat_ compared to the WT protein. ITC corroborates the KM change, with citraconate binding tighter at 20.7 μM for the Arg273Ser mutant compared to 35 μM for WT hACOD1. The ITC curve also shows saturation of the enzyme at an N value of 0.6, closer to a 1:2 binding per protein monomer than the 1:1 binding observed for WT hACOD1. As ACOD1 is a symmetric dimer, this lower N value hints at a cooperative mechanism between the active sites in the two protomers, but more work will be needed to corroborate this hypothesis. Crystal structure of hACOD1-Arg273Ser shows very clear changes in the environment around these residues. Arg273 (in the lid domain) binds to Asp417 (α-helical domain) through a single water-bridge. In contrast, Ser273 loses this tight hydrogen bonding network. Though alone this mutation does not recapitulate the increased activity of murine ACOD1, these studies indicate that it should contribute. With the active site residue Lys272 adjacent to Arg273 and the changes in hydrogen bonding network between the two domains from the point mutation and the ITC data, we suggest that this region has a dynamic allosteric role in this enzyme’s mechanism.

## 3. Conclusion

Human aconitate decarboxylase is a target of interest in the oncometabolic drug discovery community. With many studies linking itaconate production to different cancer forms, the discovery of a clinically tractable inhibitor is of great importance. To pursue structure-guided drug discovery, a detailed mechanistic and biochemical characterization is needed.

In this publication, we have expanded the biochemical, structural and dynamic knowledge of hACOD1, all aspects significantly understudied thus far. We solved the first true-apo structure of hACOD1, a necessary step towards structure-guided drug discovery, as well as the first structure of the citraconate-inhibited enzyme. We also developed a new, versatile, low-consumption, high-sensitivity, continuous kinetic assay. With the apo- and inhibited structures at hand and a robust kinetic assay showing unequivocally direct competition between the substrate (cis-aconitate) and citraconate, we confirmed the previously suggested putative active site and the mode of inhibition of this ligand.

The protein was found to have a similar structure in solution to that determined by X-ray crystallography. The dimensionless Kratky plot of the SAXS data shows no flexibility or dynamic changes between the apo and inhibited structures, indicating that any conformational changes associated with access to the active site occur on a local length-scale. Together with MD simulations and point-mutations, we demonstrated that Tyr318, though not a fully conserved, is important in both binding and catalysis and is involved in the shielding of the active site during catalysis. Local conformational changes of Tyr318 as visualized by MD are sufficient to open/close access to the small active site.

Other point mutants were also characterized. His103Ala, an active site mutation that abolishes both binding and catalysis. Arg273Ser, an allosteric mutation corresponding to the murine homologue that impacts stoichiometry of citraconate binding to the enzyme as well as the kinetic parameters. None of the point mutants showed large conformational or dynamic changes in solution. More biochemical and structural work will be needed to fully understand how point mutations in homologous proteins lead to changes in kinetic parameters as well as to determine allosteric sites for drug discovery to allow for the design of molecules beyond polydentate carboxylate mimetics.

## 4. MATERIALS AND METHODS

### 4.1. Protein expression

The expression plasmid pCAD29_hIRG1_4-461_pvp008 was a gift from Konrad Buessow (Addgene plasmid # 124843; http://n2t.net/addgene:124843; RRID:Addgene_124843)^25^. The plasmid codes for a truncated version of ACOD1, lacking short unstructured N- and C-terminal motifs. *Escherichia coli* BL21-CodonPlus (RIPL) cells (Novagen) were used for protein expression. Additionally, singe-point mutations H103A, P155A, R273S, Y318A of truncated ACOD1 (4-461) were prepared in pCOLA-Duet vector for mutational analysis of enzymatic catalysis. hACOD1 (4-461) was expressed and purified as reported previously with modifications. In brief, transformed cells were cultured in 5-10 mL of Luria-Bertani (LB) medium supplemented with 50 μg/mL kanamycin and 30 μg/mL chloramphenicol. The LB pre-culture was incubated at 37 °C with shaking at 200 rpm until the OD_600_ reached 1.2-1.5 and 1 mL of this pre-culture was used to inoculate 50 mL of LB medium. After overnight growth at 37 °C, the small LB culture was transferred to 1 L of LB medium, incubated at 37 °C with shaking at 200 rpm up to an OD_600_ of 0.75-0.80. At that point, the temperature and agitation were lowered to 22 °C and 130 rpm, respectively. After ∼1 hour, protein expression was induced with isopropylthio-β-galactoside (IPTG), at a final concentration of 500 μM. Cells were grown at 22 °C for 18 hours and harvested at 6000 x *g* for 10 min at 4 °C. Cell pellets were harvested and stored at -80 °C until further use.

### 4.2. Protein purification

Frozen cell pellets were resuspended in 50 mL of buffer A (20 mM Tris-HCl pH 8.0, 500 mM NaCl, 10% v/v glycerol, 1 mM DTT) per 20 mg of cells. The resuspension was treated with a final concentration of 1mg/mL lysozyme, 0.01 mg/mL RNAse, and 0.005 mg/mL DNAse and the cells disrupted by sonication (*Fisherbrand Qsonica 505*) at 60% power for 5 minutes (10 s pulse on and 20 s pulse off) on ice. The cellular lysate was clarified with 6-8 mL of 10% PEI followed by centrifugation at 16,500 x g for 1 hour at 4 °C. After clarification, the supernatant was loaded onto an equilibrated 5 mL Strep-Tactin XT 4Flow column (IBA Lifesciences cat#: 2-5010-025) the column washed with 5 CV of buffer A (or until UV absorbance returned to baseline) and the protein eluted in buffer A supplemented with 500 mM Biotin. The elution fractions were checked using SDS page and those containing protein were pooled and dialyzed overnight in buffer A at 4 °C with 1.5% w/w TEV protease to cleave the His_6_-tag. The protein was concentrated (3-4 mL) and further purified to homogeneity by size-exclusion chromatography (HiLoad 16/600 Superdex 200 prep grade, Cytiva) by isocratic elution with buffer B (10 mM HEPES, 150 mM NaCl, 10% v/v glycerol, 0.1 mM TCEP, pH 7.5). The purified protein was concentrated to 120 µM (6 mg/mL) and flash-frozen in small aliquots in liquid nitrogen for storage at -80 °C.

### 4.3. Protein quantification

Protein quantification was performed by UV-Vis spectroscopy, measuring the absorption at 280 nm. Accurate quantification was performed on an 8453 UV-Vis (Agilent) spectrophotometer and corrected using a three-point Morton-Stubbs correction (see supplementary information for details). The same sample was then measured on a NanoDrop™ UV-Vis Spectrophotometer (Thermo Scientific), yielding an approximate concentration, not baseline corrected. A ratio between the “true” and “approximate” concentrations was calculated and applied henceforth to all measurements routinely taken on the Nanodrop. This method showed that concentration using the Nanodrop was approximately 25% overestimated and correction yielded ITC fits for the hACOD1-citraconate binding much closer to an N of 1, validating this approach.

### 4.4. Protein crystallization screening and optimization

Initial crystallization conditions for hACOD1 were obtained from 6 sparse matrix crystallization screens (JSCG+ and Structure I+II HTS - Molecular Dimensions - and Wizard HTS, PACT HT, PEG/ION, and INDEX HT-Hampton Research). Screening was performed at room temperature (∼22 °C) using sitting drop vapor diffusion in MRC3 crystallization plates (SWISSCI) using 200 nL total volumes and varying the protein to cocktail ratio in 1:1, 2:1 and 1:2 v/v ratios for each condition and constant 40 µL reservoir volume. The drops were set using the Formulatrix NT8 drop setter, at a constant humidity of 80%. The plates were stored and imaged at 22 ºC using a Fomulatrix R1000 imager, equipped with bright field and UV absorption capabilities. From the observed protein crystals, 11 hits were identified as devoid of polydentate carboxylate components.

Two optimized conditions yielded good diffracting crystals: 1) 100 mM Tris (pH 8.8), 0.2 mM CaAc, 35% PEG4000; and B) 200 mM NaF, 35% PEG4000. Optimization was performed by varying the concentrations and pH of the crystallization cocktail components using an automated formulating system (FORMULATOR, Formulatrix) in the same tray layout described above. Co-crystallization screening of citraconate with hACOD1 was performed by varying the citraconate concentration between 2.5 mM and 10 mM to reach 99.0% occupancy based on K_d_= 35 µM measured by ITC. Fully grown crystals of both apo hACOD1 and citraconate-bound hACOD1 were obtained in 4-6 days. hACOD1 mutant proteins crystallized similarly. Drops containing crystals were layered with LV cryoOil (Mitegen) as a cryoprotectant, looped and vitrified in liquid nitrogen.

### 4.5. Data collection and reduction and structure solution

Data for the apo and citraconate-bound structures were collected at beamlines 17-ID-2 (FMX)^27^ and 17-ID-1 (AMX)^28^ in fully automated mode. Parameters for data collection can be found in Table 1. The X-ray images were indexed, integrated, scaled and merged using the automated autoproc pipeline (Global Phasing)^29^ available at NSLSII. For datasets requiring further manual data reduction (apo- and citraconate-bound hIRG1), the output of the XDS ^30^ INTEGRATE step from autoproc was parsed through pointless (within Aimless^31^, CCP4i2^32^) for space group determination and scaling. The unmerged pointless output mtz file was reduced in StarANISO^33^ (Global Phasing) to yield an anisotropic resolution cut off. Scaled and merged files from either the autoproc pipeline or from manual StarAniso runs were imported into CCP4i2, solved by molecular replacement using MOLREP^34^ with initial model PDB 6R6U (for the apo WT structure, with subsequent datasets solved using our apo structure)^25^. Rounds of manual model building using Coot^35^ and refinement using Refmac5^36^ were performed in CCP4i2. Refinement was performed using automated TLS parameters. TLS groups were used for refinement instead of anisotropic b-factors as these caused clear overfitting, with a widening gap between Rfree and Rcryst and no significant decrease in Rfree.

### 4.6. Solution NMR spectroscopy data collection and processing

1D ^1^H solution NMR spectra were collected at 298 K for cis-aconitic acid on a 14.1 T Bruker AvanceIII and 17.4 T Bruker AvanceIII spectrometer each equipped with a TCI four-channel inverse detection H/C/N/D cryoprobe. Samples were locked to 90:10 H_2_O/D_2_O. T ^1^H 1D spectra were recorded at pH 7.5 to match protein and ligand stocks. All NMR data were processed using Bruker TopSpin and MestreMnova (MestreLab Research). ^1^H chemical shifts were referenced to the water peak at 4.7 ppm. All solution spectra were analyzed using MestreMnova. ^1^H solution chemical shifts were assigned based on the published cis-aconitate and itaconate chemical shifts (Biological Magnetic Resonance Data Bank entries bmse000705 and bmse000137, respectively).

### 4.7. Enzyme Kinetic Assay

Enzyme kinetics of hACOD1 were obtained by a continuous assay observing the conversion of cis-aconitic acid to itaconic acid by ^1^H NMR. Initial velocity of enzyme progression curves was determined by measuring production of itaconate. Itaconate concentration was obtained by comparison of integrals against a standard curve of itaconate concentration.100 mM *cis*-aconitate and citraconate stock solutions were prepared using buffer B. 50-200 nM hACOD1 samples were prepared with a final *cis*-aconitate concentration ranging from 0.25-15.0 mM to a final volume of 400 µL. ^1^H 1D spectra (32K points) of final reaction mixtures were measured at 12-minute time intervals over the course of two hours at low *cis*-aconitate concentrations (0.5-3.0 mM) and 60-minute intervals for 8 hours at high *cis*-aconitate concentrations (8.0-15.0 mM). Samples prepared for citraconate inhibitory assays were measured under constant protein (50 nM) and varying substrate concentrations (20-200 µM) with citraconate concentrations ranging from 0.2 to 1.0 mM. All time points at the different conditions described were measured in triplicate.

All samples were temperature equilibrated by water-insulated heating block at 25 °C. Data was fit by Prism 10 (GraphPad) using Michaelis-Menten non-linear regression to calculate catalytic rate constant (k_cat_), maximum velocity (V_max_) and Michaelis constant (K_M_). Citraconate concentration resulting in 50% inhibition of hACOD1 activity (IC_50_) was determined by non-linear regression of sigmoidal dose-response. Each point represents three independent experiments error bars are +-1 SEM. Mode of citraconate inhibition was determine by double reciprocal plot assaying inhibitor concentration (0-1000 µM citraconate) at varying substrate concentration (1-5 mM *cis*-aconitate).

### 4.8. Small-Angle X-ray Scattering Data Collection and Analysis

In-line size-exclusion chromatography small-angle X-ray scattering (SEC-SAXS) experiments were performed at beamline 12-ID-B of the Advanced Photon Source (APS) at Argonne National Laboratory. WT apo and citraconate hACOD1 as well as single point mutants were analyzed at injection concentrations ranging from 5.0 to 6.05 mg/mL. Each sample was injected onto a Superdex 200 Increase 5/150 GL column connected to an ÄKTA micro FPLC system using a 100 μL sample loop. The samples were eluted at a flow rate of 0.3 mL/min and directed to the flow cell for simultaneous small- and wide-angle X-ray scattering (SAXS and WAXS) measurements. The running buffer consisted of 20 mM HEPES (pH 7.5), 150 mM NaCl, 0.5 mM TCEP, and 2% (v/v) glycerol. Data were collected at an X-ray energy of 13.3 keV (λ = 0.9322 Å). The combined SAXS/WAXS setup covered a momentum transfer range of 0.005 < q < 2.7 Å^−1^, where q = (4π/λ)sinθ, 2θ is the scattering angle, and λ is the X-ray wavelength. A total of 1500–2000 frames were collected per sample with an exposure time of 0.2–0.3 s per frame.

Two-dimensional scattering images were corrected for detector geometry (solid angle per pixel) and reduced to one-dimensional scattering profiles using MATLAB-based beamline software. SEC-SAXS data were further processed using BioXTAS RAW. Frames corresponding to monodisperse regions of the elution peak were identified and averaged. Background scattering was estimated from frames collected before and after the elution peak, and baseline correction was applied where necessary. The radius of gyration (Rg) was determined by Guinier analysis, using data within the range qRg≤1.3. Experimental SAXS profiles were compared with theoretical scattering curves calculated from atomic models derived from crystal structures using CRYSOL^37,38^. During fitting, parameters including the hydration layer, excluded volume, and contribution of implicit hydrogen atoms were taken into account.

### 4.9. Isothermal Titration Calorimetry

Enzyme samples for isothermal titration calorimetry (ITC) were prepared to a final concentration of 35 µM in buffer B. Citraconate was prepared to a final concentration of 1 mM by 1:1000 dilution of 1M stock in buffer B. The experiments were performed with an iTC200 Calorimeter (Malvern Panalytical/MicroCal, Netherlands/USA) at 25^°^C. The experiment consisted of 18 injections (2.1 µL each) of citraconate into a hACOD1 in the cell (200 µL) at a stirring speed of 750 RPM. A sacrificial first injection of 0.5 µL of ligand was used to accommodate the interaction during pre-titration thermal equilibration at the tip of the titration syringe; this measurement point was excluded from the final set of data. An additional set of injections was run in a separate experiment with buffer B in the cell instead of the protein solution with identical run parameters as a blank experiment for subtraction during data processing. K_d_, stoichiometry, enthalpy and entropy changes were determined from integrated binding isotherms using the “One Set of Sites” model in Origin 7.0 in the Malvern/MicroCal data analysis software.

### 4.10. MD Simulations

Simulations were performed in GROMACS^39^. Each was started from apo-hACOD1 (residues 4-461) solvated in a rhombic dodecahedral water box; charges were neutralized with sodium and chloride ions and addition 150 mM NaCl was present to reproduce experimental conditions. The system was equilibrated for 1 ns in the NPT ensemble before starting production runs. Ten replicates of 200 ns with a 2 fs time step were run in parallel starting from the same equilibrated structure. Trajectories were analyzed with MDAnalysis ^40^. The distance between the Cα of Tyr318 and Cα of Pro155 was measured for each protomer over every frame in the trajectories. The Cα root mean square fluctuations (RMSF) were determined by first calculating an average structure over all trajectories, aligning to this structure and calculation of the RMSF using the average structure as a reference.

## Acknowledgements

We thank Daniel McVicar and Jonathan Weiss (Cancer Innovation Laboratory, NCI, NIH) for fruitful discussions and an introduction to the world of cancer immunology. Thank you to Asokan Anbanandam and the NMR Facility for Biological Research for assistance with data acquisition of NMR experiments and to Marzena Dyba within the CCR Biophysics Resource for access to instrumentation and experimental help and training. This work also utilized the computational resources of the NIH HPC Biowulf cluster (https://hpc.nih.gov).

We thank the NCI SAXS core facility for support in collection and analysis of SAXS data in-house and at beamline 12-ID-B at the Advanced Photon Source on beam time award through PUP1008087 proposal. NCI SAXS Facility are supported by the Frederick National Laboratory for cancer research (contract 75N91019D00024) and the NIH Intramural Research Program. This research also used resources 17-ID-1 and 17-ID-2 of the National Synchrotron Light Source II.

