## Supplementary information for "A robust structural, kinetic and biophysical characterization of human ACOD1 to support novel chemotherapeutic development"

Supplemental information

Sup. Table 1 hACOD1 WT and mutant expression information

Sup. Table 2 List of screening crystallization conditions

Sup. Table 3 hACOD1 Crystallization conditions

Sup. Fig. 1 ^1^H NMR spectrum of inhibitory assay of hACOD1

Sup. Fig. 2 SAXS profiles of hACOD1 mutants

Sup. Fig. 3 Baseline correction of protein UV-Vis absorbance Spectra

Sup. Table 1 hACOD1 WT and mutant expression information

| Source |  |
| --- | --- |
| Source organism | Homo sapiens |
| DNA source | synthetic |
| Expression vector | pCAD29_hIRG1_4-461_pvp008[1] (addgene #124843) |
| Expression host | *E. coli* |
| Expression details | Heterologous protein expression of hACOD1 was carried out in CodonPlus (RIPL) BL21. 1 L LB cultures were inoculated to a final IPTG concentration of 500 [µM] for 18 hours at 22°C and 130 rpm agitation. |
| Complete amino-acid sequence of the WT protein produced, with point mutants highlighted as:  His103Ala – Yellow  Arg273Ser – Pink  Tyr318Ala - Green | MASWSHPQFEKVDENLYFQ­GGGRKSITESFATAIHGLKVGHLTDRVIQRSKRMILDTLGAGFLGTTTEVFHIASQYSKIYSSNISSTVWGQPDIRLPPTYAAFVNGVAIHSMDFDDTWHPATHPSGAVLPVLTALAEALPRSPKFSGLDLLLAFNVGIEVQGRLLHFAKEANDMPKRFHPPSVVGTLGSAAAASKFLGLSSTKCREALAIAVSHAGAPMANAATQTKPLHIGNAAKHGIEAAFLAMLGLQGNKQVLDLEAGFGAFYANYSPKVLPSIASYSWLLDQQDVAFKRFPAHLSTHWVADAAASVRKHLVAERALLPTDYIKRIVLRIPNVQYVNRPFPVSEHEARHSFQYVACAMLLDGGITVPSFHECQINRPQVRELLSKVELEYPPDNLPSFNILYCEISVTLKDGATFTDRSDTFYGHWRKPLSQEDLEEKFRANASKMLSWDTVESLIKIVKNLEDLEDCSVLTTLLKGP |

Sup. Table 2 List of crystallization conditions that produced three-dimensional crystals at least 25 µm in size which lacked additives structural homologous to substrate. All conditions were screened for optimal pH, [salt], and [PEG].

| **Crystallization Screen** | **Well #** | **Additive** | **Buffer** | **Precipitant** |
| --- | --- | --- | --- | --- |
| Index | F2 | TMAO [0.2M] | Tris pH 8.0 [0.1M] | PEG 2000 [20%w/v] |
| Index | G4 | LiSO_4_ [0.2M] | HEPES [0.1M] | PEG 3350 [25%w/v] |
| JCSG+ | G4 | TMAO [0.2M] | Tris pH 8.5 [0.1M] | PEG 2000 [20%w/v] |
| PACT | D7 | NaCl [0.2M] | Tris pH 8.0 [0.1M] | PEG 6000 [20%w/v] |
| PACT | D11 | CaCl_2_ [0.2M] | TRIS pH 8.0 [0.1M] | PEG 6000 [20%w/v] |
| PACT | F1 | NaF [0.2M] | BIS-TRIS propane pH 6.5 [0.1M] | PEG 3350 [20%w/v] |
| PACT | F3 | NaI [0.2M] | BIS-TRIS propane pH 6.5 [0.1M] | PEG 3350 [20%w/v] |
| PACT | F5 | NaNO_3_ [0.2M] | BIS-TRIS propane pH 6.5 [0.1M] | PEG 3350 [20%w/v] |
| PACT | F6 | Na formate [0.2M] | BIS-TRIS propane pH 6.5 [0.1M] | PEG 3350 [20%w/v] |
| Structure | C9 | MgCl_2_ [0.2M] | TRIS pH 8.5 [0.1M] | PEG 4000  [30% w/v] |
| Structure | D11 | NaOAc [0.2M] | Tris pH 8.5 [0.1M] | PEG 4000  [30% w/v] |

Sup. Table 3 hACOD1 Crystallization conditions

| Protein form | Citraconate-bound (9O5N) | Apo (9O5J) | Y318A | H103A | R273S |
| --- | --- | --- | --- | --- | --- |
| Method | Sitting drop | | | | |
| Plate type | Swissci MRC3 | | | | |
| Temperature (°C) | 20 ºC | | | | |
| Protein concentration | 5.1 mg/mL | 5.9 mg/mL | | | |
| Ligand concentration | 15 mM | N/A | | | |
| Buffer composition of protein solution | 10 mM HEPES pH 7.5, 150 mM NaCl, 10% glycerol, 1 mM TCEP | | | | |
| Composition of reservoir solution | 100 mM Tris pH 8.0, 25% PEG 4000, 200 mM CaOAc | 100 mM Tris pH 8.8, 35% PEG 4000, 200 mM CaOAc | | | |
| Volume and ratio of drop | 200 nL, 2:1 protein:reservoir | | | | |
| Volume of reservoir | 40 uL | | | | |
| Composition of the cryoprotectant | Mitegen LV CryoOil | | | | |
| Drop setting | Formulatrix Formulator (reservoir) and NT8 (drop setting) | | | | |
| Seeding | No | | | | |

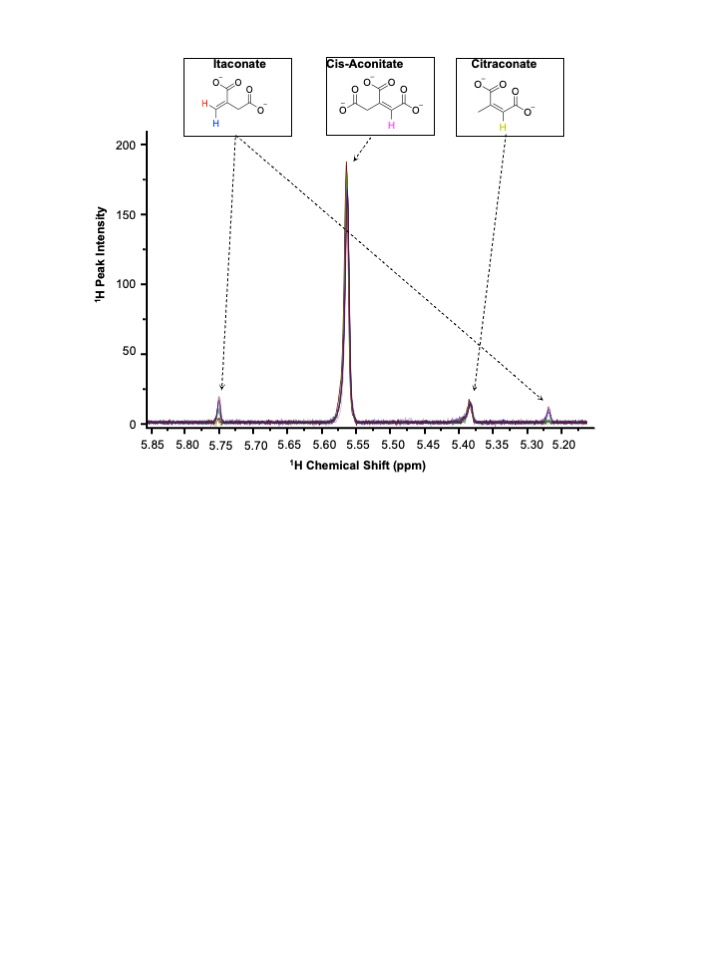

Sup. Fig. 1 ^1^H NMR spectrum of inhibitory assay of hACOD1 by dose-dependence of citraconate, showing the 1H peaks associated with the three different carboxylate species.

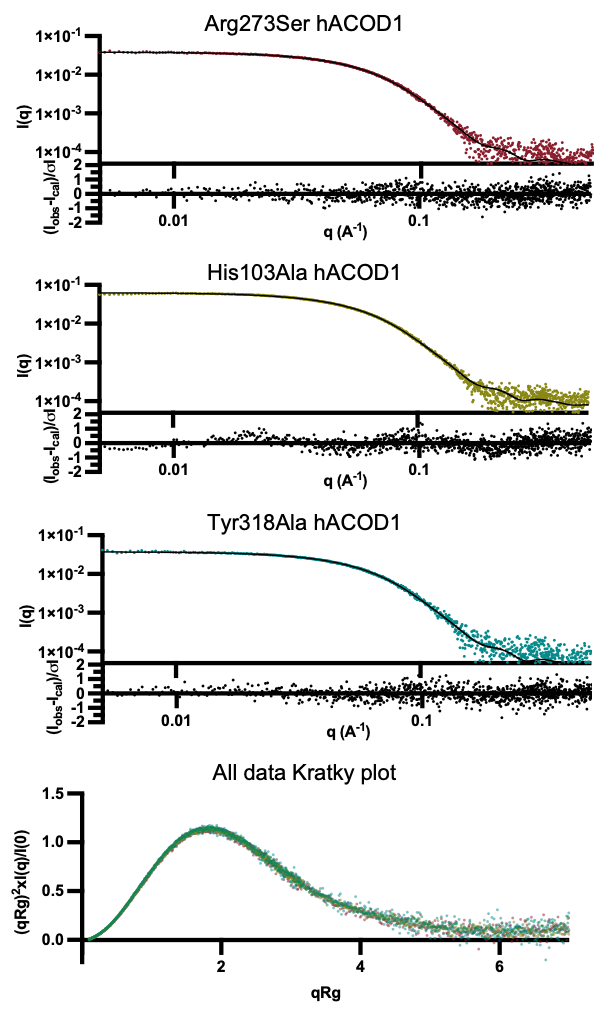

Sup. Fig. 2 SAXS profiles of hACOD1 mutants fitted to the calculated solution scattering profiles derived from their corresponding X-ray crystal structures, along with dimensionless Kratky plots for all datasets. hACOD1 WT in green, citraconate-bound in blue, Arg273Ser in red, His103Ala in yellow and Tyr318Ala in teal. The dimensionless Kratky plot shows low dynamics, similar across all datasets

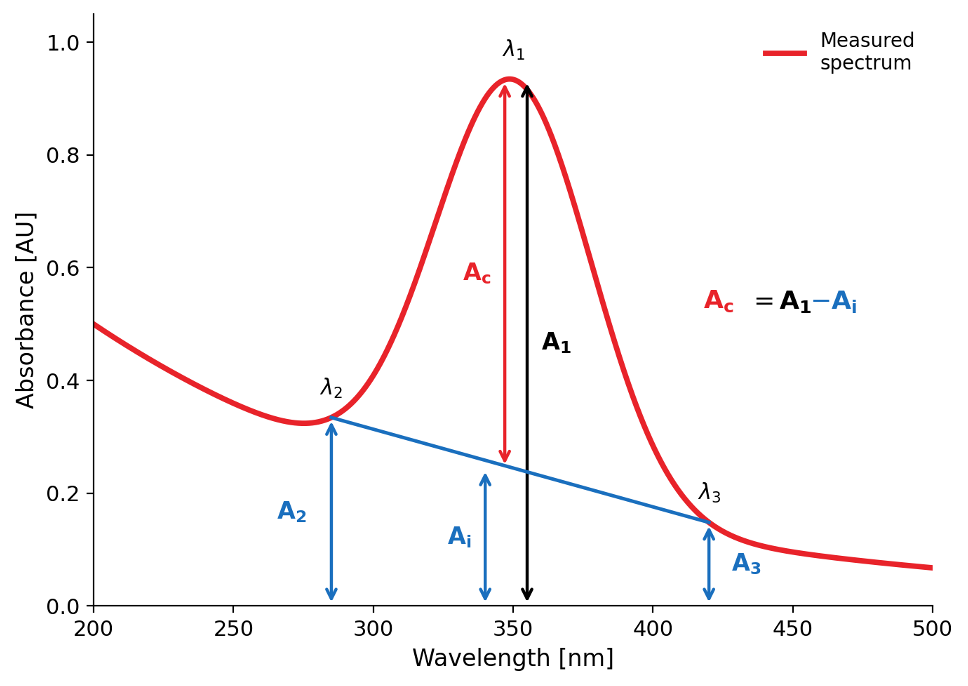

Sup. Fig. 3 Baseline correction of protein UV-Vis absorbance Spectra. Representative UV-Vis absorbance spectrum of a protein sample (red curve) showing a peak at λ_1_ (~280 nm). A linear baseline (blue line), defined between reference wavelengths λ_2_ and λ_3_  was used to estimate the background absorbance (A_i_) at λ_1_. The corrected absorbance (A_c_) was calculated as A_c_ = A_1_ – A_i_, where A_i_ is the measured absorbance at λ_1_. Arrows indicate the raw absorbance (A_1_), and corrected absorbance (A_c_).

Equation 1.1 $A_{i}=\frac{\left( A_{2}-A_{3} \right)}{2}$

Equation 1.2 $A_{c}\left( corrected OD \right)= A_{1}-A_{i}$

Measurements were performed in a QS high-precision quarts cuvette (Hellma Analytics) with a 10 mm length. Samples (70µL each) were analyzed following blank correction with Buffer B. Prior to measurements, the cuvette was cleaned by rinsing with deionized water, washing with 3% (v/v) Hellmanex™ III solution, and thoroughly rinsing again with deionized water before air drying. Absorbance was recorded at 280 nm, and protein concentrations were calculated using a three-point correction.
